# SARS-CoV-2 spike S2 subunit inhibits p53 activation of p21(WAF1), TRAIL Death Receptor DR5 and MDM2 proteins in cancer cells

**DOI:** 10.1101/2024.04.12.589252

**Authors:** Shengliang Zhang, Wafik S. El-Deiry

**Affiliations:** Laboratory of Translational Oncology and Experimental Cancer Therapeutics, Warren Alpert Medical School, Brown University, Providence, Rhode Island, USA; Department of Pathology and Laboratory Medicine, Warren Alpert Medical School, Brown University, Providence, Rhode Island, USA; Joint Program in Cancer Biology, Lifespan Health System and Brown University, Providence, Rhode Island, USA; Legorreta Cancer Center at Brown University, Providence, Rhode Island, USA; Hematology/Oncology Division, Department of Medicine, Lifespan Health System and Brown University, Providence, Rhode Island, USA

**Keywords:** SARS-COV2 spike, p53, MDM2, chemotherapy, cancer

## Abstract

Severe acute respiratory syndrome coronavirus 2 (SARS-CoV-2) and COVID-19 infection has led to worsened outcomes for patients with cancer. SARS-CoV-2 spike protein mediates host cell infection and cell-cell fusion that causes stabilization of tumor suppressor p53 protein. In-silico analysis previously suggested that SARS-CoV-2 spike interacts with p53 directly but this putative interaction has not been demonstrated in cells. We examined the interaction between SARS-CoV-2 spike, p53 and MDM2 (E3 ligase, which mediates p53 degradation) in cancer cells using an immunoprecipitation assay. We observed that SARS-CoV-2 spike protein interrupts p53-MDM2 protein interaction but did not detect SARS-CoV-2 spike bound with p53 protein in the cancer cells. We further observed that SARS-CoV-2 spike suppresses p53 transcriptional activity in cancer cells including after nutlin exposure of wild-type p53-, spike S2-expressing tumor cells and inhibits chemotherapy-induced p53 gene activation of p21(WAF1), TRAIL Death Receptor DR5 and MDM2. The suppressive effect of SARS-CoV-2 spike on p53-dependent gene activation provides a potential molecular mechanism by which SARS-CoV-2 infection may impact tumorigenesis, tumor progression and chemotherapy sensitivity. In fact, cisplatin-treated tumor cells expressing spike S2 were found to have increased cell viability as compared to control cells. Further observations on γ-H2AX expression in spike S2-expressing cells treated with cisplatin may indicate altered DNA damage sensing in the DNA damage response pathway. The preliminary observations reported here warrant further studies to unravel the impact of SARS-CoV-2 and its various encoded proteins including spike on pathways of tumorigenesis and response to cancer therapeutics.

## Introduction

Severe acute respiratory syndrome coronavirus 2 (SARS-CoV-2) has led to more severe outcomes of patients afflicted with cancer [1]. Cancer development has been a controversial area as a potential long-term effect of SARS-CoV-2 infection [2]. Previous studies have shown that SARS-CoV-2 proteins could increase breast and lung cancer cell proliferation, migration and invasion [3, 4]. There is a need to investigate and better understand whether SARS-CoV-2 may be involved in any way with cancer signaling pathways, molecular mechanisms of cancer development or has effects on therapy.

SARS-CoV-2 spike protein is a key mediator for virus infection of host cells though its two subunits: S1, which binds to human angiotensin-converting enzyme 2 (ACE2), and S2, which mediates a membrane fusion process [5]. The S2 subunit contains multiple domains that facilitate protein-protein interaction. Therefore, the S2 subunit is a factor to explore functional effects of SARS-CoV-2 spike protein in host cells after virus entry.

An in-silico analysis using HADDOCK 2.2 software previously suggested that p53 and BRCA1/2 may interact with the heptic repeat-2 region of the S2 subunit through C-terminal domain [6]. DNA damage or therapy-induced tumor suppressor p53 protein transcriptionally activates genes leading to multiple effects preserving genome integrity, altering metabolism, immune response, cell cycle, DNA repair, cell growth and cell apoptosis to prevent or eliminate transformed cells [7]. Loss of p53 function increases the incidence of carcinogen-induced tumorigenesis and drives chemo-resistance [8]. SARS-CoV-2 infection has been found to alter p53 stabilization. The previous studies have shown that SARS-CoV-2 spike in particular plays a role to stabilize and activate p53 by mediating cell-cell fusion or induction of ROS in host cells during SARS-CoV-2 virus infection [9, 10]. In response to cellular stress, activated p53 regulates specific gene expression, including MDM2 (E3 ligase). MDM2, in turn, binds to p53 and triggers p53 ubiquitination and proteasomal degradation [11], while interruption of MDM2-p53 interaction leads to p53 stabilization. Thus, a putative interaction between SARS-CoV-2 spike, p53 and p53 related signaling pathways following SARS-CoV-2 infection could impact cellular homeostasis, tumorigenic pathways, and/or response to cancer therapeutics.

In this study, we performed cell-based assays to examine the effect of SARS-CoV-2 on p53 activation in cancer cells and demonstrate that SARS-CoV-2 spike interrupts the MDM2-p53 interaction in cancer cells and alters p53 signaling in cancer cells upon chemotherapy included blunted activation of p53 targets involved in growth arrest and apoptosis.

## Methods

### Cell culture

Human cell lines used in this study include human lung cancer cells H460 (ATCC), breast cancer cells MCF7 (ATCC), colorectal cancer cells HCT116 (p53 wild-type or p53-null) and sarcoma cells U2OS with p53-knockout (U2OS-P53KO). HCT116 and U2OS-P53KO cells were cultured in McCoy’s 5A (modified) medium, while H460 cells were cultured in RPMI-1640 medium and MCF7 cells were cultured in Eagle’s Minimum Essential Medium. All cell line media were supplemented with 10% FBS and 1% Penicillin-Streptomycin. The cell lines were authenticated and tested to ensure the cultures were free of mycoplasma infection. H460 (lung cancer), MCF7 (breast cancer), HCT116 (colon cancer), and U2OS (osteosarcoma) are the most commonly studied wild-type p53-expressing human tumor cell lines across tumor types.

### Plasmid transfection

The plasmids pcDNA3.1-SARS2-spike (#145032) and p-CMV-Neo-Bam-p53wt (#16434) were obtained from Addgene. The plasmids were transiently transfected with lipofectamine 2000 (Life Technologies, catalog no. 11668-027) into cancer cells as stated in the figures.

### Luciferase reporter assay

Cancer cells were transfected with an equal amount of each plasmid as indicated in the figures. The PG13-luciferase reporter expression in cancer cells was examined based on bioluminescence using the IVIS imaging system (PerkinElmer, Hopkin, MA, USA) at different time points as indicated in the figures.

### Western blot analysis

Cells were seeded with the same density on culture plates and were lysed in loading buffer (Sigma-Aldrich, St. Louis, MO, USA). Equal amounts of cell lysates were electrophoresed through 4-12% SDS-PAGE, then transferred to PVDF membranes. The transferred PVDF membranes were blocked with 5% skim milk at room temperature, then incubated with primary antibodies incubated in a blocking buffer at 4°C overnight. Antibody binding was detected on PVDF with appropriate HRP-conjugated secondary antibodies by a Syngene PXi imaging system (Syngene). Anti-sars-spike antibodies (NB100-56578) were purchased from NOVUS Biologicals, and anti-p53 (DO-1, catalog no. sc-126), anti-MDM2 (SMP14, catalog no. sc-965), and anti-p53 (FL-393) antibodies were purchased from Santa Cruz Biotechnology. Anti-p21 (Ab-1, catalog no. OP64) and anti-Noxa (catalog no. OP180) antibodies were purchased from EMD Millipore. Anti-cleaved PARP (catalog no. 9546) and γ-H2AX (catalog no. 2577) antibodies were purchased from Cell Signaling Technology.

### Immunoprecipitation

Cell lysates were incubated with 2 μg of anti-p53 antibody (DO-1) overnight at 4°C; then, they were mixed with 50 μL of nProtein A Sepharose 4 Fast Flow (Cytiva, catalog no. 17528001) for 3 hours at 4°C and washed with lysis buffer three times. The immunoprecipitated proteins were eluted from the nProtein A-Sepharose beads by boiling with 2× sample buffer (Invitrogen, catalog no. NP0007) and subjected to SDS-PAGE.

### CellTiter-Glo luminescent Cell viability assay

Cell viability was measured by CellTiter-Glo bioluminescence (Promega, catalog no. G7572), and analyzed using an IVIS imager.

### Statistical analysis

The statistical significance of differences between pairs was determined using Student’s t tests with GraphPad Prism. The minimal level of significance was P < 0.05.

## Results

### SARS-CoV2-spike overexpression shows reduced p53 interaction with MDM2 in cancer cells

To investigate the interaction between SARS-CoV-2 spike, p53, and MDM2 proteins in cancer cells, we performed an immunoprecipitation assay. The pcDNA3.1-SARS2-spike plasmid was introduced to overexpress SARS-CoV-2 spike protein containing the S2 subunit in cancer cells [12]. The plasmids p-CMV-Neo-Bam-p53wt, pcDNA3.1-SARS2-spike and p-EGFP-MDM2 were co-transfected into p53-knockout U2OS (U2OS-p53KO) cancer cells using lipofectamine.

The immunoprecipitation assay showed that MDM2 protein bound with p53 in the cells while cells with SARS-CoV-2 spike S2 subunit overexpression displayed reduced amounts of MDM2 bound with p53 when compared to the pcDNA3.1 transfection control (**Figure 1A**). These results suggest that SARS-CoV-2 spike S2 subunit overexpression can alter p53 binding with MDM2 in cancer cells.

**Figure 1.**
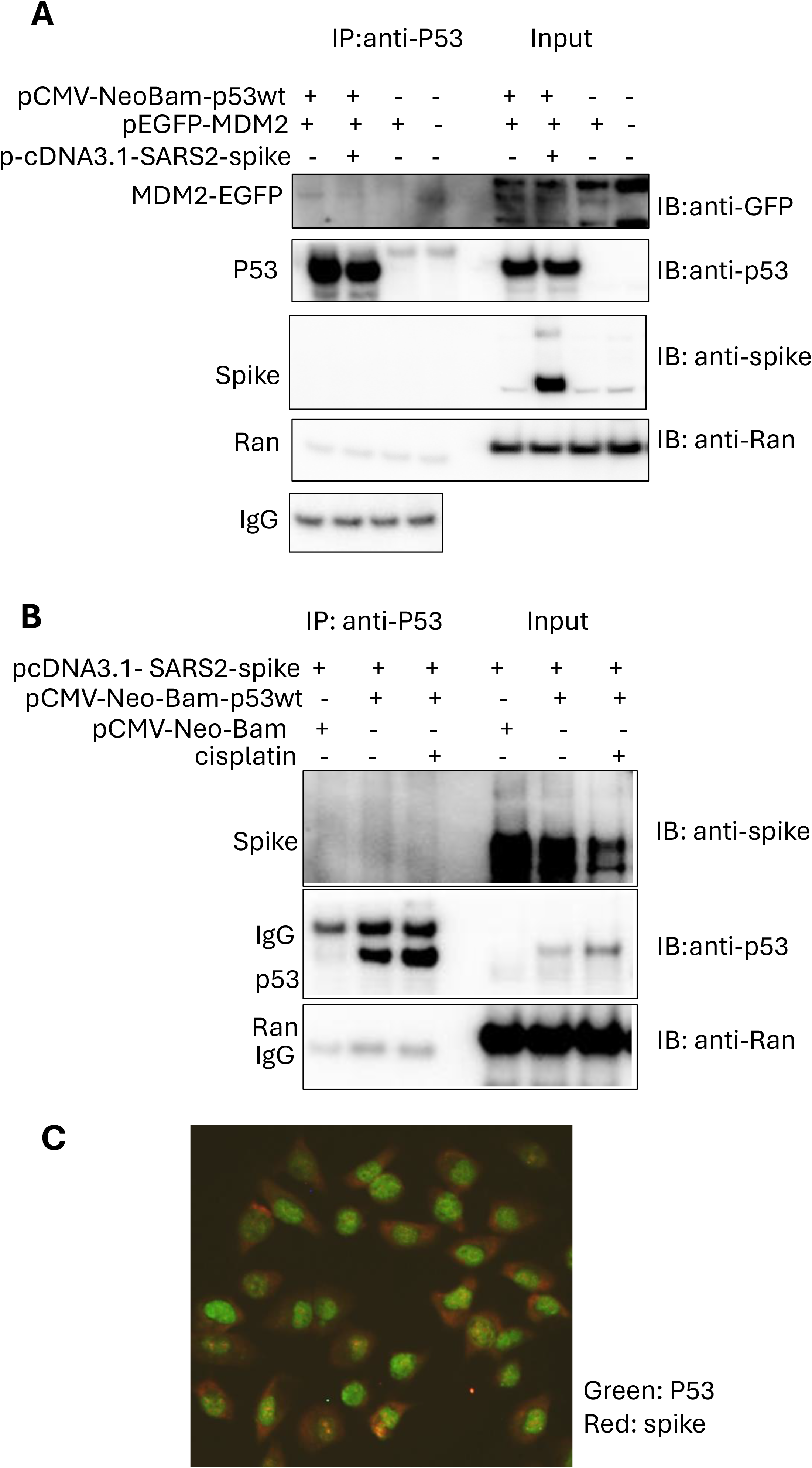
Reduced interaction between p53 and MDM2 following SARS-CoV-2 spike S2 overexpression in cancer cells. A. Immunoprecipitation (IP) assay in U2OS-p53KO cancer cells. Cells were transiently transfected with the plasmids as indicated. IP was performed with anti-p53 (DO-1) and IB with anti-p53 (FL393). B. Immunoprecipitation in p53 wild-type breast cancer cells. MCF7 cancer cells were treated with cisplatin for 4 hours. IP was performed with anti-p53 (DO-1). C. Immunofluorescence imaging the cellular locations of the SARS-CoV-2 spike S2 and p53. H460 cancer cells were treated with 3 μM of cisplatin for 24 hours.

However, SARS-CoV-2 spike S2 subunit was not observed to bind with p53 protein in the immunoprecipitation assay (**Figure 1A**), nor did it have any detectable impact when p53 was activated by treatment with cisplatin, a DNA damaging agent that causes interstrand crosslinks (**Figure 1B**).

Our observations from lack of co-immunoprecipitation between p53 and the SARS-CoV-2 spike S2 protein subunit are consistent with different cellular locations of SARS-CoV-2 spike S2 and p53 in the cancer cells treated with cisplatin (**Figure 1C**). The immunofluorescence imaging showed that the majority of p53 was localized in the nuclei, while the majority of SARS-CoV-2 spike S2 subunit was localized in the cytoplasm in H460 cells treated with cisplatin (**Figure 1C**). These results do not demonstrate SARS-CoV-2-spike S2 subunit protein binding to wild-type p53 in cancer cells either in the absence or presence of cisplatin treatment. As this is a protein subunit, and some small amount of nuclear staining is observed, we cannot exclude that spike S2 can gain access to the nucleus or that intact spike could do so as well.

### SARS-CoV2 spike attenuates p53 transcriptional activity in cancer cells

To investigate the effect of the SARS-CoV2 spike S2 subunit on p53 signaling in cancer cells further, we conducted a PG13-luciferase (PG13-Luc) reporter assay. PG13-luc contains 13 copies of the p53-binding consensus sequence, and the production of PG13-luc was confirmed by using bioluminescence in cells [13]. The p53-null HCT116 and U2OS-p53KO cancer cells were transiently transfected with PG13-luc together with p-CMV-Neo-Bam-p53wt and pcDNA3.1-SARS2-spike for 20 hours. The cells with pcDNA3.1-SARS2-spike transfection showed reduction of the p53 responsive bioluminescence, as compared to the pcDNA3.1 transfection control (**Figure 2A**). Further treatment with nutlin-3a, an MDM2 inhibitor which activates p53 signaling, was ineffective at rescuing the reduction of the p53 responsive bioluminescence of PG13-Luc (**Figure 2B**).

**Figure 2.**
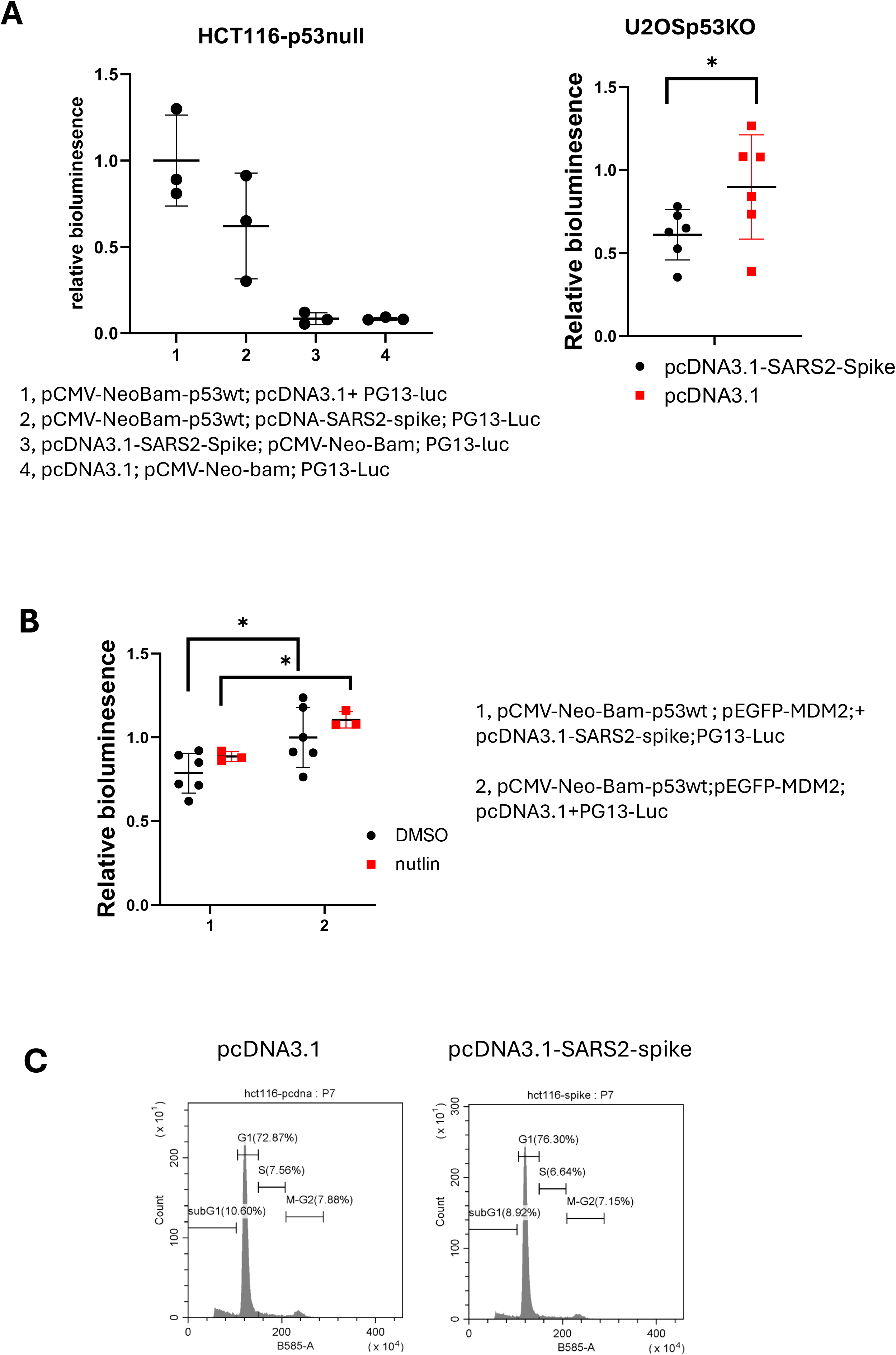
SARS-CoV-2 spike S2 suppresses p53-mediated transcriptional activity in cancer cells. A. PG13-luc reporter assay in cancer cells. HCT116 p53-null cells and U2OS-p53KO were transiently transfected with p-CMV-Neo-Bam-p53wt and PG13-luc together with the plasmids as indicated for 20 hours. B. PG13-luc reporter assay. U2OS-p53KO cells were transfected with the plasmids as indicated, followed with 10 μM of nutlin-3a treatment for 4 hours. C. Cell cycle profiling in HCT116 cells transiently transfected with the plasmids as indicated for 72 hours.

The suppressive effect of spike on p53 responsive bioluminescence was also detected in a variety of p53 wild-type cancer cell lines by either transient transfection or stable expression of PG-13Luc (**Figures 3A and 3B**). These results suggest that SARS-CoV-2 spike S2 subunit protein attenuates p53 transcriptional activity. In a preliminary experiment following up on the inhibition of p53 activity by SARS-CoV-2 spike S2 subunit protein, no cell cycle arrest was detected at G1, S or G2-M phases in cancer cells transfected with pcDNA3.1-SARS2-spike, as compared to the pcDNA3.1 transfection control (**Figure 2C**).

**Figure 3.**
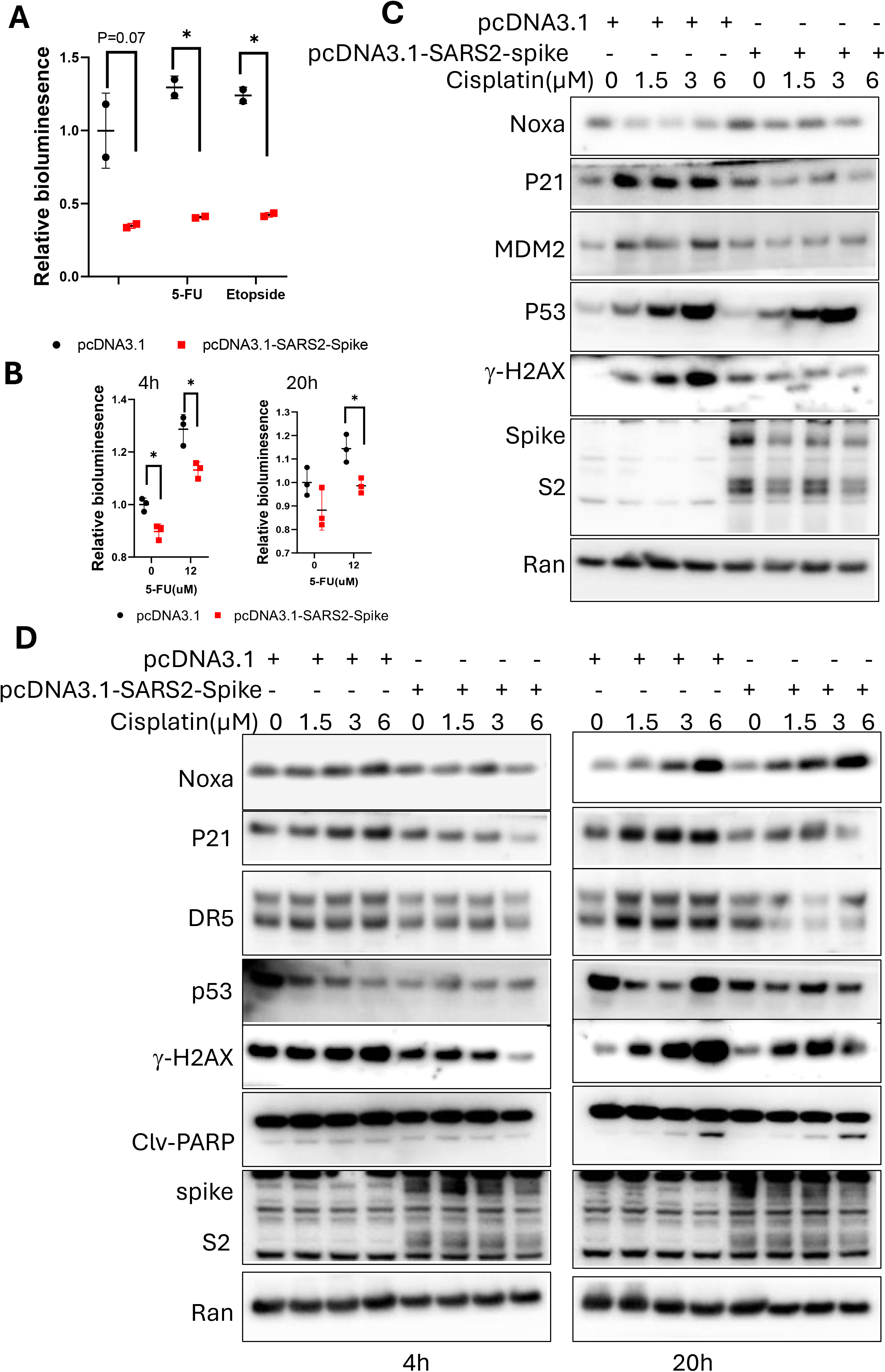
SARS-CoV-2 spike S2 abrogates chemotherapy-induced p53-mediated activation of p21(WAF1), TRAIL Death Receptor DR5 and MDM2 in cancer cells. A. PG13-luc reporter assay in cancer cells. Breast cancer MCF7 cells were transiently transfected with pcDNA3.1-SARS2-spike S2 and PG13-luc, followed by exposure to 100 μM of etoposide or 200 μM of 5-FU for 20 hours. B. PG13-luc reporter assay in HCT116-PG13-luc stable cells. HCT116-P13-Luc cells were transiently transfected with the pcDNA-SARS2-spike S2, followed by 12 μM 5-FU treatment for 4 and 20 hours. C. Protein analysis of p53 targets in MCF7 cells. The cells were transiently transfected with pcDNA3.1-SARS2-spike S2, followed by cisplatin treatment for 20 hours. D. Protein analysis of the p53 targets in H460 cells. The cells were transiently transfected with pcDNA3.1-SARS2-spike S2, followed by cisplatin treatment for 4 and 20 hours.

### SARS-CoV-2 spike S2 subunit reduces p53 unpregulation of p21(WAF1) and TRAIL Death Receptor DR5 proteins as well as γ-H2AX levels after cisplatin treatment in cancer cells

We further investigated if the SARS-CoV-2 spike S2 subunit protein can inhibit chemotherapy-induced p53 transcriptional activity in cancer cells. p53 wild-type breast cancer cells MCF7 and lung cancer cells H460 were transiently transfected with the pcDNA3.1-SARS2-spike S2 subunit and PG13-luc, followed by etoposide or 5-Fluorouracil (5-FU) treatment. The PG13-luciferase reporter assay showed a reduction of the p53-responsive bioluminescence in the cells transfected with pcDNA3.1-SARS2-spike S2 subunit, as compared to the pcDNA3.1 transfection control (**Figure 3A**). A similar reduction of p53-responsive transcription-mediated bioluminescence was also observed in p53 wild-type HCT116-PG13-luc cells which have been stably transduced with PG13-luc (**Figure 3B**). These results suggest that SARS-CoV-2 spike S2 subunit reduces chemotherapy-induced p53 transcriptional activity in cancer cells.

We further examined endogenous p53 targets at the protein level in cancer cells upon cisplatin treatment. Consistent with the reduction of the p53 responsive bioluminescence, a decrease or delay in the p53 transcriptional targets, p21, TRAIL Death Receptor DR5 and MDM2 at the protein level was detected in cancer cells transfected with the pcDNA-SARS2-spike S2 subunit, as compared to the pcDNA3.1 transfection at different post-treatment time points (**Figures 3C and 3D**). These results suggest that SARS-CoV-2-spike alters chemotherapy-induced p53 signaling in cancer cells of pathways involved in growth arrest and cell death.

p53 is involved in DNA damage response as well as repair [14]. We examined the DNA damage repair response by analyzing γ-H2AX level in cancer cells. The cisplatin treatment caused an increase of γ-H2AX at the protein level in the cancer cells (**Figure 3C and 3D**), indicating DNA damage repair response in p53 wild-type cancer cells. The levels of the γ-H2AX were reduced in the cohort of the cells transfected with pcDNA3.1-SARS2-spike, as compared to the pcDNA3.1 transfection control (**Figure 3C and 3D**). These results suggest that the SARS-CoV-2 spike causes an altered DNA damage sensing and repair response in cancer cells.

We further examined the effect of SARS-CoV-2 spike S2 subunit protein on cell growth and death in cancer cells upon chemotherapy treatment. H460 lung cancer cells were transiently transfected with pcDNA3.1-SARS2-spike S2 subunit, followed by cisplatin treatment for 40 hours. The cisplatin treatment increased the PARP cleavage, a death marker, in the H460 cells with pcDNA3.1 transfection at the tested doses (**Figure 4A**). A similar PARP cleavage was also observed in H460 cells transfected with pcDNA3.1-SARS2-spike S2 subunit. An increase in cell viability was observed in H460 cells transfected with pcDNA-SARS-CoV-2 spike (**Figure 4B**) at the tested doses of cisplatin, as compared to pcDNA-3.1 transfection.

**Figure 4.**
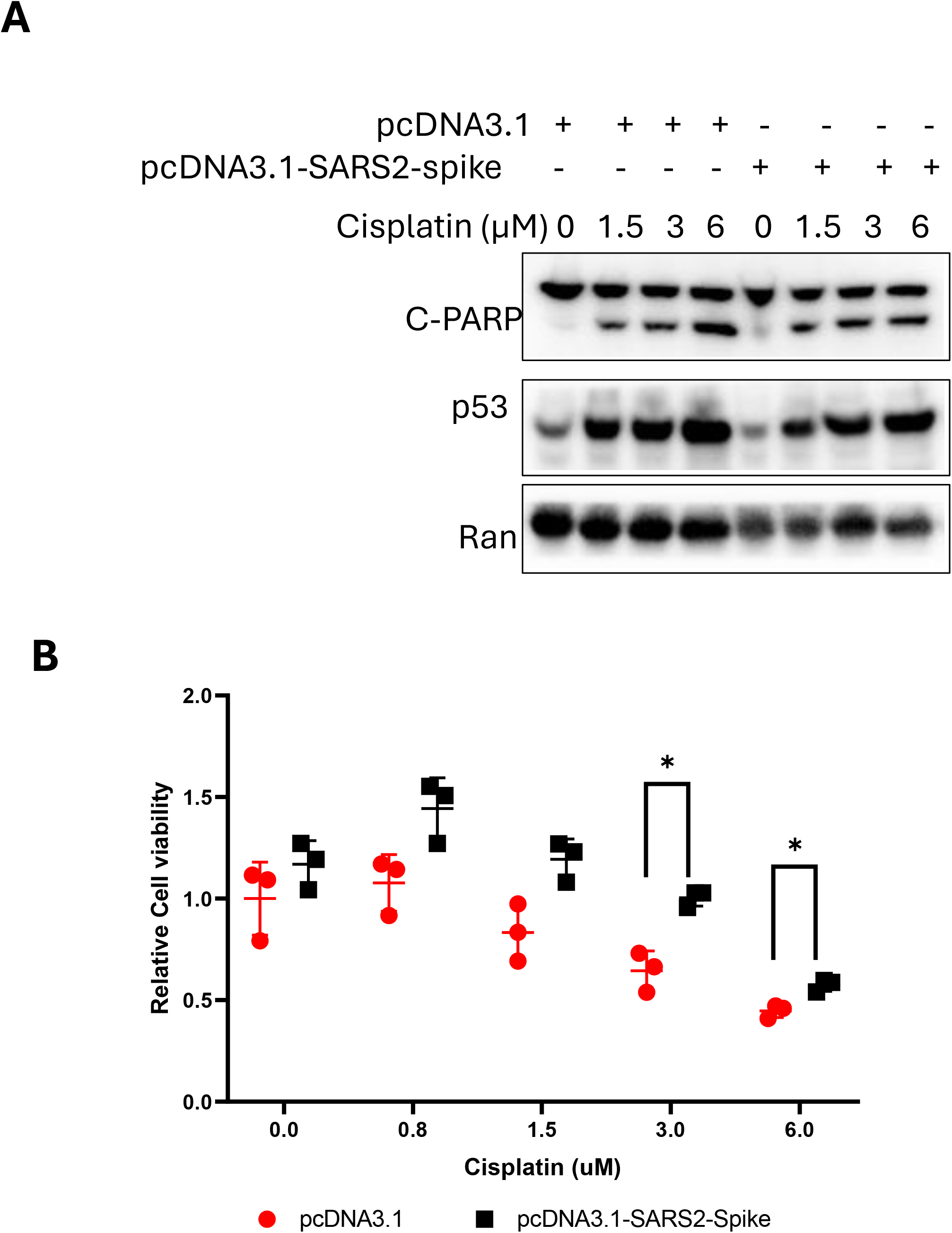
The effect of SARS-CoV-2 spike S2 on cell growth and death. The cancer cells were transiently transfected with pcDNA-SARS2-spike S2, followed by cisplatin treatment. A. Protein analysis of PARP cleavage in H460 cancer cells. The cells were transfected with pcDNA-SARS2-spike S2, followed with cisplatin treatment for 40 hours. B. Cell viability assay. H460 cells were transfected with pcDNA-SARS2-spike S2, followed with cisplatin treatment for 72 hours.

## Discussion

The SARS-CoV-2 spike S2 protein subunit plays a key role in SARS-CoV-2 invasion of human cells through spike S2 binding to receptor angiotensin-converting enzyme 2 (ACE2) on the human host cell surface. We show here that spike S2 can alter p53 transcriptional activity in wild-type p53-expressing cancer cells based on reduction of the p53-responsive reporter activity and a decrease in selected p53 targets such as p21(WAF1) or TRAIL Death Receptor DR5 at the protein level.

Our findings differ from previous reports that have shown that SARS-CoV-2 spike stabilized p53 and activated p53 [9, 10]. In the previous studies, the p53 activation and stabilization was caused by the spike-ACE2 mediated cell-cell fusion and an increase in ROS in cancer or normal cells [9, 10].

To study the effect of spike S2 on p53 activity as would occur after virus entry into host cells, we transiently transfected SARS-CoV-2 spike S2 subunit into cancer cells thereby avoiding possible effects of a virus infection or cell-cell fusion on p53 signaling. Our findings agree with previous results showing that spike could not stabilize p53 in low ACE2 cells that are unable to undergo cell-cell fusion [9].

The p53 protein is considered as a “genome guardian” by arresting the cell cycle to repair DNA damage or causing cell death in the presence of unrepaired persistent damage and stress [14]. In our preliminary experiments reported here, the SARS-CoV-2 spike-induced inactivation of p53 was correlated to an apparent reduction of expression of DNA damage response protein γ-H2AX after cisplatin exposure and a reduced cell cycle checkpoint response in the cancer cells (**Figure 3**). Whether these changes are a consequence from the suppressive effect of SARS-CoV-2 spike S2 on p53 signaling in cancer cells needs to be further investigated.

SARS-CoV-2 spike has been found to regulate multiple signaling pathways [15]. The post-translational modifications in p53 including phosphorylation and/or acetylation generally result in activation of p53 through different signaling pathways [16]. It remains unclear whether SARS-CoV-2 spike S2 interrupts p53 binding to the DNA promoters of the targets, or whether SARS-CoV-2 spike inhibits p53 transcriptional activity through post-translational modifications in p53 or other alterations in proteins that complex with p53.

We found that SARS-CoV-2 spike S2 interrupts p53-MDM2 interaction in cancer cells in the absence of exposure to DNA damaging agents. These results were observed by an immunoprecipitation assay using exogenously overexpressed p53 and MDM2 in cancer cells. The effect of SARS-CoV-2 spike S2 on endogenous p53 binding to MDM2 needs to be investigated in more detail in the future. Nevertheless, our observation provides a possible molecular mechanism by which SARS-CoV-2 spike S2 mediates p53 stabilization caused by cell-cell fusion.

We did not observe SARS-CoV-2 spike S2 binding with p53 in cancer cells using immunoprecipitation assay as previously predicted through an in-silico study showing that the S2 subunit of SARS-CoV-2 spike may interact with p53 [6]. In our study, we performed an immunoprecipitation assay in two cancer cell lines transiently transfected with the plasmids carrying SARS-CoV-2 spike S2. The cell conditions in this study are not the same as SARS-CoV-2 infections that cause syncytia formation. SARS-CoV-2 causes severe disease in multiple organs, and severe effects on cell function that might confer specific post-translational modifications (PTMs). Specific PTMs on spike might modulate host factor binding [17]. It is possible that spike protein with some potential PTMs might interact with p53 under some cellular conditions that were not simulated in our experiments. We note and have no current understanding of the increased γ-H2AX observed on western blot at basal conditions with spike S2 in MCF7 in absence of cisplatin treatment (**Figure 3C**). This increased basal γ-H2AX expression in spike S2-expressing cells was not observed in H460 cells at two different time-points (**Figure 3D**). Future studies will need to determine the veracity of the MCF7 result and its meaning if correct.

While spike S2 appeared to attenuate the induction of p21(WAF1), TRAIL Death Receptor DR5 and MDM2 after DNA damage, there was less of an apparent effect on pro-apoptotic Noxa in our preliminary studies. However, under the experimental conditions, we did not observe more PARP cleavage at the time points evaluated. With cisplatin treatment there was greater cell viability when spike S2 was overexpressed. Additional work needs to determine long-term effects on cell viability, effects on cell survival proteins in chemotherapy-treated spike S2 or SARS-CoV-2 infected cells, in addition to more detailed studies of the DNA damage and repair effects of spike S2. For the latter studies, host cell reactivation assays could be performed with reporters treated with cisplatin either in a tube before transfection or following transfection of human cells with or without spike S2.

We have not conducted in vivo experiments and some of our experiments lack additional controls such as in flow analysis or by looking at kinetics of cell cycle checkpoint regulation. We have not evaluated normal cells such as airway, muscle, immune, brain or intestinal cells. Cycling vs quiescent cells are also important to investigate for potential differential effects of spike or other SARS-CoV-2 proteins. We have not investigated immune cell interactions such as NK or T-cells in our experiments where spike S2 protein was overexpressed in culture. These would all be reasonable early future directions.

Our results have implications for the biological effects of spike S2 subunit in human cells whether spike is present due to primary COVID-19 infection or due to mRNA vaccines where its expression is used to promote anti-viral immunity. A perturbed p53 pathway is concerning but also complicated in sorting out since cellular transformation and cancer are a multi-step process that evolves over time. Further detailed studies can more fully characterize the effects of spike, as well as structural determinants within the protein for interaction between the DNA damage sensing and response pathways as well as the p53 tumor suppressing pathway. With respect to the p53 pathway, further studies are needed to unravel how less MDM2 is bound to p53 in the presence of spike and the mechanisms underlying reduced p21(WAF10), TRAIL Death Receptor DR5 as well as MDM2 under conditions where there is less degradation of p53 due to reduced interaction with MDM2.

P21(WAF1) and TRAIL Death Receptor DR5 levels often go up in stress and through many pathways but here we see lack of induction. Why the p53 response is blunted in the presence of spike S2 remains an open question. The effect of spike S2 on γ-H2AX can be interpreted in a different way too although one would not expect either less damage or accelerated DNA repair with cisplatin. Our interpretation is reduced DNA damage sensing and response after cisplatin exposure in the presence of spike S2 but that remains to be further unraveled. One can investigate the ordered events in the DNA damage sensing and response in different cellular backgrounds including repair-deficient cells such as BRCA mutation or other cancer susceptibility states (ATM, mismatch repair, Fanconi, PTEN, Wnt/beta-catenin/APC, Rb, etc.). Future studies can investigate effects of spike on other cancer pathways, oncogenic or tumor suppressive as well as at a broader range of therapeutic efficacies isn the presence of spike or other SARS-CoV-2 encoded proteins. As there is already a history of various viruses associated with human cancer including hepatitis viruses HBV/HCV, EBV, HPV, and potentially SV40, SARS-CoV-2 is a candidate that should be further investigated.

In summary, we identified the SARS-CoV-2 spike S2 subunit as a COVID-19 virus factor that interrupts p53 binding to MDM2 in cancer cells and demonstrated the suppressive effect of SARS-CoV-2 spike S2 on p53 signaling in cancer cells. Correlated to the inhibition of p53 signaling, the short-term expression of spike S2 caused an altered DNA damage response through altered levels of γ-H2AX after DNA damage in cells, altered sensing in the damage response to cisplatin Importantly, the p53-dependent DNA damage induction of growth arrest and apoptotic targets p21(WAF1) and TRAIL Death Receptor DR5 was significantly attenuated under different experimental conditions with spike S2 and this was associated with greater cell viability in the presence of spike S2 and chemotherapy treatment. As loss of p53 function is a known driver of cancer development and confers chemo-resistance, our study provides insight into cellular mechanisms by which SARS-CoV-2 spike S2 may be involved in reducing barriers to tumorigenesis during and post SARS-CoV-2 infections.

## Acknowledgements

W.S.E-D. is an American Cancer Society Research Professor and is supported by the Mencoff Family University Professorship at Brown University. This work began early during the COVID pandemic when it was supported by a Brown University pilot grant. Subsequent experiments were made possible by discretionary start-up funds to Dr. El-Deiry at Brown University.

